# Bacterial community studies and novel *Bradyrhizobium* and *Rhizobium* strains from drought-tolerant legumes grown in Namibia

**DOI:** 10.1101/2025.08.11.667748

**Authors:** Paidamoyo N. Mataranyika, Cristina Bez, Alessio Mengoni, Francesca Vaccaro, Oluwaseyi S. Olanrewaju, Percy M. Chimwamurombe, Jean Damascene Uzabakiriho, Vittorio Venturi

**Affiliations:** Department of Life Sciences, Silwood Park Campus, Imperial College London, Ascot, United Kingdom; International Centre for Genetic Engineering and Biotechnology, Trieste, Italy; Department of Biology, University of Florence, Florence, Italy; Unit for Environmental Sciences and Management, Microbiology, North-West University, Private Bag X6001, Potchefstroom, 2520, South Africa; Department of Biology, Chemistry and Physics, School of Natural and Applied Sciences, Namibia University of Science and Technology, Windhoek, Namibia; Department of Biochemistry, Microbiology and Biotechnology, University of Namibia, Windhoek, Namibia; African Genome Center, University Mohammed VI Polytechnic (UM6P), Ben Guerir, Morocco

**Keywords:** Rhizobia, Plant Microbiome, Nodule Endophytes, Drought-tolerant Legumes, Abiotic Stress, Namibia

## Abstract

**Aims:** Analysis of the microbiota associated with root nodules of five species of drought-tolerant legumes grown in Namibia. These legumes were *Lablab purpureus, Vigna radiata, Vigna unguiculata, Macrotyloma uniflorum,* and *Vigna aconitifolia*.

**Methods:** 16S rRNA gene amplicon sequencing analysis and isolation of rhizobial and non-rhizobial bacterial strains from root nodules was performed. Plant growth-promoting traits were assessed on some isolates and the genomes of four rhizobial strains were sequenced.

**Results:** The microbiota analysis revealed *Bradyrhizobium* as the most prevalent genus in the root nodules, while most of the non-rhizobial community included members of *Bacillus* genus. In addition, a strain of *Variovorax paradoxus* was also isolated from *V. unguiculata*, representing the first documented isolation and characterization report in Namibia. *In vitro* phenotypic characterization of the non-rhizobial nodule endophytes indicated that they possessed several plant growth-promoting traits. The four rhizobial strains isolated were taxonomically affiliated to the genera *Bradyrhizobium* and *Rhizobium*. Genome sequencing revealed that these strains possibly belong to novel species since they share only a limited similarity (dDDH values<70%) to already known species.

**Impact Statement:** This study demonstrated that drought-tolerant legume species harbour microbial diversity from Namibian soil, which allowed us to increase our knowledge on plant-associated bacteria and their possible use in sustainable farming systems, as potential bioinoculants for crop production and for soil revitalization. Utilizing locally isolated bacterial strains as microbial bioinoculants maximizes their compatibility with the local conditions increasing the potential for positive effects on plant health and ecosystem functioning.

## Introduction

Diazotrophic bacteria, which can fix atmospheric nitrogen, play a vital and sustainable role when in association with legumes, by contributing nearly 65% of biologically fixed nitrogen in the agricultural industry (1). Particularly relevant are symbiotic nitrogen-fixing bacteria, known as rhizobia. They colonise roots of leguminous plant species giving rise to the formation of specialized organs, called nodules, where biological nitrogen fixation (BNF) take place (2). This offers alleviation of nitrogen deficiencies in soils in subsistence farmin in Africa (3).

Rhizobia are found in several genera of the phylum Pseudomonadota, particularly in the classes of *Alphaproteobacteria* and *Betaproteobacteria*. Examples are *Cupriavidus* (4), *Bradyrhizobium* (5)*, Rhizobium* (6)*, Mesorhizobium* and *Sinorhizobium* (7). Strains from these genera can harbour nitrogen fixing and nodulating genes allowing root nodule formation and BNF (8). Nodules can also be colonized by non-rhizobial bacterial species (9), which can behave like commensal endophytes and in some cases can also affect (positively or negatively) the rhizobial species (10).

There is an emerging interest in developing biofertilizers and/or biocontrol agents based on plant growth-promoting bacteria (PGPB). This is due to their plant growth promoting traits including, but not limited to, nitrogen fixation (11), phosphate solubilisation (12), 1-aminocyclopropane-1-carboxylate (ACC) deaminase production (13), and direct as well as indirect biocontrol activities towards plant pathogens (14). The exploitation of these PGPB as biofertilizers provides an alternative environmentally friendly and sustainable way for improving soil fertility which is essential for crop production (15).

To date, there are limited resources and collections on plant microbiota in Namibia and in the entire African continent (16). Given the growing challenges related to food security and the increasing costs of agricultural inputs, there is a pressing requirement to explore more environmentally and economically sustainable alternatives to fertilizers (17–19) such as the exploitation of the local beneficial plant microbiota (20).

Moreover, it is important to use locally isolated bacteria as bioinoculants, as these have evolved and have acclimatized to specific local environmental factors. Indeed, the exploration and exploitation of the plant-associated microbial biodiversity can offer opportunities to develop novel biotechnological solutions, translate bioinoculant formulations on local crop species, and contribute to biodiversity (21,22). Consequently, in order to fully harness the potential of local microbial diversity in legume species cultivated in Namibia, it is essential to begin to establish a collection of microbes associated with these legumes.

This work presents culture independent and dependent studies of root nodule endophytes of drought-tolerant legumes grown in Namibia. To the best of our knowledge, it is also the first report of studies related to the *Bradyrhizobium* genus of the root nodule microbiota of a set of drought tolerant legumes grown in Namibia. The legumes investigated are dolichos [*Lablab purpureus* (L.) Sweet var. Lignosus Prain], mung bean [*Vigna radiata* (L.) R. Wilczek var. radiata], cowpea (*Vigna unguiculata* L. Walp), horsegram (*Macrotyloma uniflorum* Var. Madhu), and mothbean [*Vigna aconitifolia* (Jacq.) Marechal].

## Materials and methods

### Seed material and growth conditions

This study was done in Namibia at the University of Namibia using six accessions from five legume species obtained from National Bureau of Plant Genetic Resources (NBPGR), India in July 2019. These were [IC0623025 (*L. purpureus*), Gujarat 5 (*V. unguiculata*), Himala (*M. uniflorum*), IC39399 (*V. radiata*), and two accessions from *V. aconitifolia*: IPCMO-880 and RMB-25]. Microbiota studies were carried out on four legume accessions-IC0623025 (*L. purpureus*), Gujarat 5 (*V. unguiculata*), Himala (*M. uniflorum*), and IPCMO-880 (*V. aconitifolia*). Rhizobia were isolated from IC0623025 (*L. purpureus*), and Gujarat 5 (*V. unguiculata*). Four rhizobial strains from IC0623025 (*L. purpureus*) and Gujarat 5 (*V. unguiculata*) were sequenced.

Legumes were grown in pots in a greenhouse between February and March with soil obtained from Bagani, Kavango East Region, Namibia. Seeds (four) were planted in triplicates of eight pots with soil obtained from Bagani, Kavango East. Samples were grown in pots in a greenhouse at the University of Namibia, Windhoek main campus. Atmospheric humidity in the greenhouses was maintained at 25% with natural light hours during February and March. The plants were watered twice a week, each receiving 200 ml of water. Plants were grown for 5 weeks before harvesting root nodules.

The Kavango East region experiences a hot semi-arid climate. Temperatures average around 23°C with normal temperatures exceeding 30°C most of the year and a minimum of 10°C in the winter seasons. Annual rainfall ranges between 500 mm and 700 mm (23). The soil type is dominantly arenosol, however, small patches of calcisol and solonetz are present in the Kavango East region. The soil pH is typically 7.50, with total nitrogen, 0.06%, organic carbon 0.48%, phosphorous 58.20 ppm^1^, potassium 0.90 me^2^ %, calcium 1.30 me^2^ %, magnesium 1.70 me^2^ %, manganese 0.18 me^2^ %, copper 0.60 ppm^1^, iron 0.70 ppm^1^, zinc 0.50 ppm^1^, and sodium 0.09 % (24).

### 16S rRNA amplicon library preparation and sequencing

DNA was extracted from approximately 100mg of surface-sterilised nodules from a single plant [as previously described (25)] using QIAGEN^®^ DNeasy^®^ Plant Mini Kit (Qiagen, USA, Valencia, CA) following the manufacturer’s instructions. To obtain the required nodule material (by mass), nodules from the same plant were pooled in cases where more than one nodule was required. Samples from *V. aconitifolia* (RMB-25) and *V. radiata* (IC0623025) were excluded as nodule mass could not be attained for DNA extraction. Surface sterilisation was confirmed by inoculating 100µL of the final sterilisation wash onto sterile MAG medium plates and incubated for 7 days at 30°C (25). Samples that showed microbial growth after 7 days were excluded from further analysis. Microbiota library preparation was done following the Illumina MiSeq System Manual. The protocol targeted the 16S V3 and V4 region using the primers Forward Primer = 5’ (314F) TCGTCGGCAGCGTCAGATGTGTATAAGAGACAGCCTACGGGNGGCWGCAG and Reverse Primer = 5’ GTCTCGTGGGCTCGGAGATGTGTATAAGAGACAGGACTACHVGGGTATCTAAT CC (805R) (26), and Illumina Whole Genome Sequencing was performed by SeqCentre LLC (Pittsburgh, PA). Sequence reads were submitted to GenBank (under BioProject PRJNA896769) and assigned accession numbers are shown in Supplementary Table 2. Rhizobia were isolated using modified arabinose gluconate (MAG) that supports the growth of nitrogen-fixing rhizobia. Macerate from surface-sterilised nodules was inoculated onto MAG plates incubated at 30°C for 10 days. Pure cultures were maintained on MAG (27).

### Amplicon data processing

The .fastq files were imported into QIIME2 version 2022.8 (28), the clustering of reads into Amplicon Sequence Variants ASVs was done using the DADA2 plugin (29) and taxonomic assignment was done based on SILVA database (release 138) (30). The dataset was imported in R using the package qiime2R (31), and the subsequent analyses and plots were drawn using either phyloseq or microbiome R-packages (32,33) and visualized by using ggplot package in R software (34). The prevalence of each taxon was determined at the level of ASV and Genus. Community richness and diversity were determined using the QIIME diversity core-metrics-phylogenetic command for alpha and beta diversity analysis in the QIIME2 package. We estimated the Chao1 and Shannon diversity (Hʹ) OTU richness indices using the Vegan (35) and Phyloseq packages (32) in R. Bray-Curtis distance was calculated from the normalized ASV tables using the function ordinate of the R package Vegan (35).

### Whole genome sequencing

Strains identified as rhizobia were further analysed based on their genome sequences. The strains were grown in modified arabinose gluconate (MAG) broth at 30°C for 7days. DNA was subsequently extracted using the Norgen Biotek Corp. Bacterial Genomic DNA Isolation Kit. The complete genomes of *L. purpureus* and *V. unguiculata* nodule bacterial endophytes were sequenced with the Illumina MiSeq platform using 150 bp paired-end reads and following the tagmentation Illumina Nextera XT protocol (Illumina Inc., San Diego, CA, USA). Paired-end reads from each isolate were quality controlled using FastQC (v.0.11.7) (36) and adapted and trimmed using Trimmomatic v.0.39 (37). Trimmed raw data were further assembled de novo using SPAdes (v3.15.5) (38) algorithm to create a draft genome sequence for each isolate. To evaluate the genome assembly quality Quast (v.5.0.2) (39) was used and completeness and contamination was assessed by using a set of genes that are ubiquitous and present in single copy within a phylogenetic lineage with CheckM (v.1.1.6). Kraken2 (v.2.0.8) was used as a taxonomic sequence classifier to assign taxonomic labels to DNA sequences. The assembled genomes were annotated, using the NCBI Prokaryotic Genome Annotation Pipeline (PGAP). Genomes were also annotated using RAST (Rapid Annotation using Subsystem Technology) Server (40), uploaded to the Integrated Microbial Genomes and Metagenomes (IMG/M) database and automatically annotated, using annotation pipeline IMG Annotation Pipeline v.4.16.6 (41). Functional annotation was performed by DFAST (42) ran from https://dfast.ddbj.nig.ac.jp/.

### Taxonomic identification

Genome-based phylogeny and digital DNA-DNA hybridization [dDDH, (43)] were computed to taxonomically place the strains by the Type (Strain) Genome Server (TYGS) (44), using default options. Phylogenetic inference was done on TYGS web server and dDDH clustering was done using a 79% dDDH threshold as previously introduced (45). Pairwise comparisons among the 4 genomes were also performed by running JSpecies (46), which allows computation of ANI (based on BLAST+) and tetranucleotide comparisons.

Trees inferred with FastME 2.1.6.1 (47) from Genome BLAST Distance Phylogeny (GBDP) distances calculated from genome sequences. The branch lengths are scaled in terms of GBDP distance formula *d5*. The numbers above the branches are GBDP pseudo-bootstrap support values > 60 % from 100 replications, with an average branch support from 77.3% to 90.4% (see **Supplementary Table 5**). The tree was rooted at the midpoint (48).

Secondary metabolites gene clusters have been identified by AntiSMASH (49) (accessed on 10/04/2024). To provide genome-based indication of the biosafety issues of selected strains, antibiotic resistance determinants were inspected from PGAP annotated genomes. Protein sequences were extracted from Genbank (.gb) files using custom Python scripts and used as input for the analyses. Antibiotic resistance determinants have been identified by Comprehensive Antibiotic Resistance Database CARD database search (50) accessed on 2024.04.18. (51)

Search for *nodC* and *nifH* orthologs were done by performing local BLAST search (NCBI BLAST+ blastp) using annotated genomes on the Galaxy web server (https://usegalaxy.eu/). Annotations files from DFAST (52), PGAP (53), and KAAS (54) were used.

### Isolation and characterization of nodule endophytes

Nodules were harvested, surface sterilised, and prepared for isolation following a similar method previously described (55). Surface sterilisation was confirmed by inoculating 100µL of the final sterilisation wash onto sterile yeast extract mannitol (YEM) medium plates and incubated for 7 days at 30°C. Samples that showed microbial growth after 7 days were excluded from further analysis. Thereafter, 100µL of each macerated sample was inoculated onto YEM and modified soil extract agar (SEA) plates (56). Plates were incubated at room temperature for 3 days before being transferred to a 30°C incubation chamber. Growth was monitored over a period of 5-7 days. Pure cultures were maintained on YEM and modified SEA plates.

### Molecular characterization of bacterial nodule isolates

Isolates were identified following a method similarly described by Pesce, Kleiner & Tisa, (57) using 16S rRNA universal primers: FDIFuni-5′-AGA GTT TGA TCC TGG CTC-3′ and P2Runi-5′-ACG GCT ACC TTG TTA GGA CTT-3′ and sequenced with primers 907r (5′-CCGTCAATTCMTTTRAGTTT-3′) (58) and F785 (5′-GGATTAGATA-CCCTGGTA-3′) (59). Samples were sequenced by Eurofins Genomics, Germany. Primary sequence data was run through the National Centre for Biotechnology Information Basic Local Alignment Search Tool (NCBI BLAST) to determine identity (60). Accession numbers obtained after submission onto GenBank fall under project PRJNA896769 are shown in Supplementary Table 1.

### Evaluation of PGP activity of bacterial nodule endophytes

Phosphate solubilization (61), siderophore production (62), exopolysaccharide (EPS) production (63), indole acetic acid (IAA) production (64), and heavy metal tolerance (65) were assessed as previously described. Isolates were grown on Jensen medium (66) as an indication for biological nitrogen fixation. Additional molecular tests were carried out for rhizobia. Antifungal activity against *Fusarium graminearum,* was tested following the method described by Rajendran et al. (67). The results of the bioactivity of the isolates are detailed in Supplementary Table 1 and summarised in Figure 3. Effect of rhizobia on nodulation was also determined on *V. unguiculata* (cowpea seeds) as previously described (11). Cowpea seeds were surface sterilised and incubated in PBS solutions with respective strains. This experiment included seeds that were not inoculated which represented the control. Nodulation was monitored over 30 days.

## Results

### Culture-independent microbiota analysis of root nodules

To evaluate the representativeness of cultured isolates and the presence of additional taxa in the root nodules (viz. the overall root microbiota diversity), a 16S rRNA gene amplicon sequencing analysis was performed from DNA purified from nodules. The amplicon sequences were cleaned to exclude non-bacterial reads and low sequence reads; no ASVs were identified as Archaea and reads annotated as chloroplast and mitochondria were excluded from the data set. After these exclusions, a total of 17364 reads were further analysed. Average reads per legume species ranged from 3171 to 5677 with *L. purpureus* having the highest number of reads.

Not surprisingly, the genus *Bradyrhizobium* was very abundant in all samples (Figure 1B); on average, it made up 99.2% and 98% of all bacterial reads in *L. purpureus* and *M. uniflorum* nodules respectively. In *V. unguiculata,* 72.9% were *Bradyrhizobium* while *V. aconitifolia* (IPCMO-880) had the least amount of *Bradyrhizobium* reads with an average of 69.9% detected.

**Figure 1:**
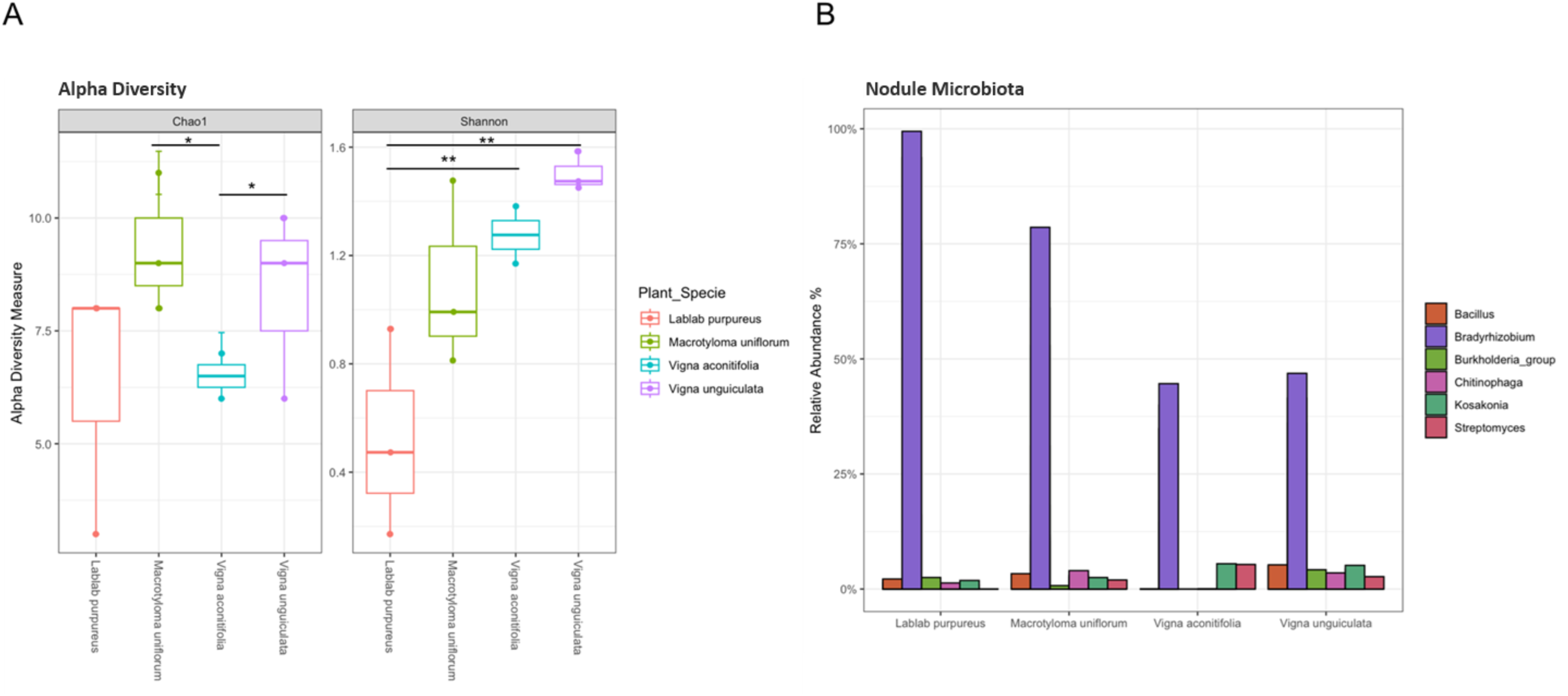
Diversity of root nodule microbiota. A) Chao1 and Shannon diversity values, represented as boxplots colored according to the four species of legumes grown in Namibia and used in this study. Significant diversity at the Wilcoxon test is defined as *p<0.05 and **p<0.01); B) Bar chart showing the average relative abundance of the most abundant taxa detected in the nodule microbiota in each legume species. Bars are colored according to the bacteria genus. In Figure 1B, relative abundances were recalculated by filtering for the taxa of interest within each sample category and rescaling their total to 100%.

Reads assigned to other bacterial genera were also detected. The genus *Bacillus* and its taxonomic relatives accounted for most of these bacteria and reads belonging to *Streptomyces* and *Xanthomonas* spp. were also identified. The alpha diversity indices (Shannon and Chao1), as presented in Figure 1A, showed significant differences in the community biodiversity of the four legumes species, suggesting that the legume genotypes could have played an important role in the recruitment of the bacterial community in the conditions tested here. In particular, *L. purpureus* and *M. uniflorum* exhibited minimal variation, whereas *M. uniflorum* and *V. aconitifolia* (IPCMO-880) were most similar in the number of observed taxa. However, in general, the biodiversity indexes show very low values suggesting that the nodules were a very close/intimate and specific plant niche.

### Isolation of rhizobial strains from legume root nodules

Four rhizobial strains were isolated from root nodules of *L. purpureus* and *V. unguiculata* named hereafter as CB7, CB9, DN3 and DN5. These isolates belonged to *Bradyrhizobium* (isolated from *L. purpureus*) and *Rhizobium* (isolated from *V. unguiculata*) genera (see below, assigned accession numbers are listed in Supplementary Table 1).

### Whole genome sequencing of the four rhizobial strains

In order to obtain more information on the isolated rhizobial strains, their genome was sequenced. The genomes features are provided in Table 1. Taxonomic assignment was performed by comparing genome sequences with type strain sequences present in TYGS. Results of dDDH values span from 19.9% to 22.4% for CB7, from 19.9% to 22.3% for CB9, from 28.9% for DN3 and DN5, significantly lower than the 70% threshold, indicating that these strains could represent novel species. Additional tables reporting the details of the type strains used for comparisons are reported in Supplementary material SM1.

**Table 1.** Summary of functional annotation (DFAST).

|  | CB7 | CB9 | DN3 | DN5 |
| --- | --- | --- | --- | --- |
| Total Seq Length (bp) | 7633888 | 7641318 | 8454736 | 8455917 |
| N° of Contigs | 104 | 151 | 121 | 129 |
| Longest Seqs (bp) | 899202 | 933950 | 715429 | 846548 |
| **N50 (bp)** | 339507 | 290806 | 229046 | 313285 |
| **GC content (%)** | 59.6 | 59.6 | 63.2 | 63.2 |
| **N°of CDSs** | 7295 | 7298 | 7881 | 7940 |
| **gene** | <sup>1</sup><br>— | <sup>1</sup><br>— | <sup>1</sup><br>— | <sup>1</sup><br>— |
| **Average Protein Length** | 301.2 | 301.1 | 297.2 | 295.9 |
| **Coding Ratio (%)** | 86.4 | 86.3 | 83.1 | 83.3 |
| **rRNAs** | 3 | 3 | 3 | 3 |
| **tRNAs** | 49 | 49 | 54 | 54 |
| **tmRNA** | <sup>1</sup><br>— | <sup>1</sup><br>— | <sup>1</sup><br>— | <sup>1</sup><br>— |
| **CRISPRs** | 0 | 0 | 0 | 0 |

<sup>1</sup>Dashes indicate that the feature is not listed in the output.

Phylogenetic trees based on WGS (Figure 2) indicated that DN3 and DN5 were affiliated to *Bradyrhizobium*, while CB7 and CB9 to the genus *Rhizobium*. These trees also supported that the strains could be novel species, since they do not cluster with known reference strains (see Supplementary Material SM1 for details on clustering with type strains). Pairwise ANI comparison between CB7 and CB9, and DN3 and DN5 indicated that they are the same species and possibly very closely related strains (Supplementary Table 5) since the results in values are higher than 99.8 ANI%.

**Figure 2.**
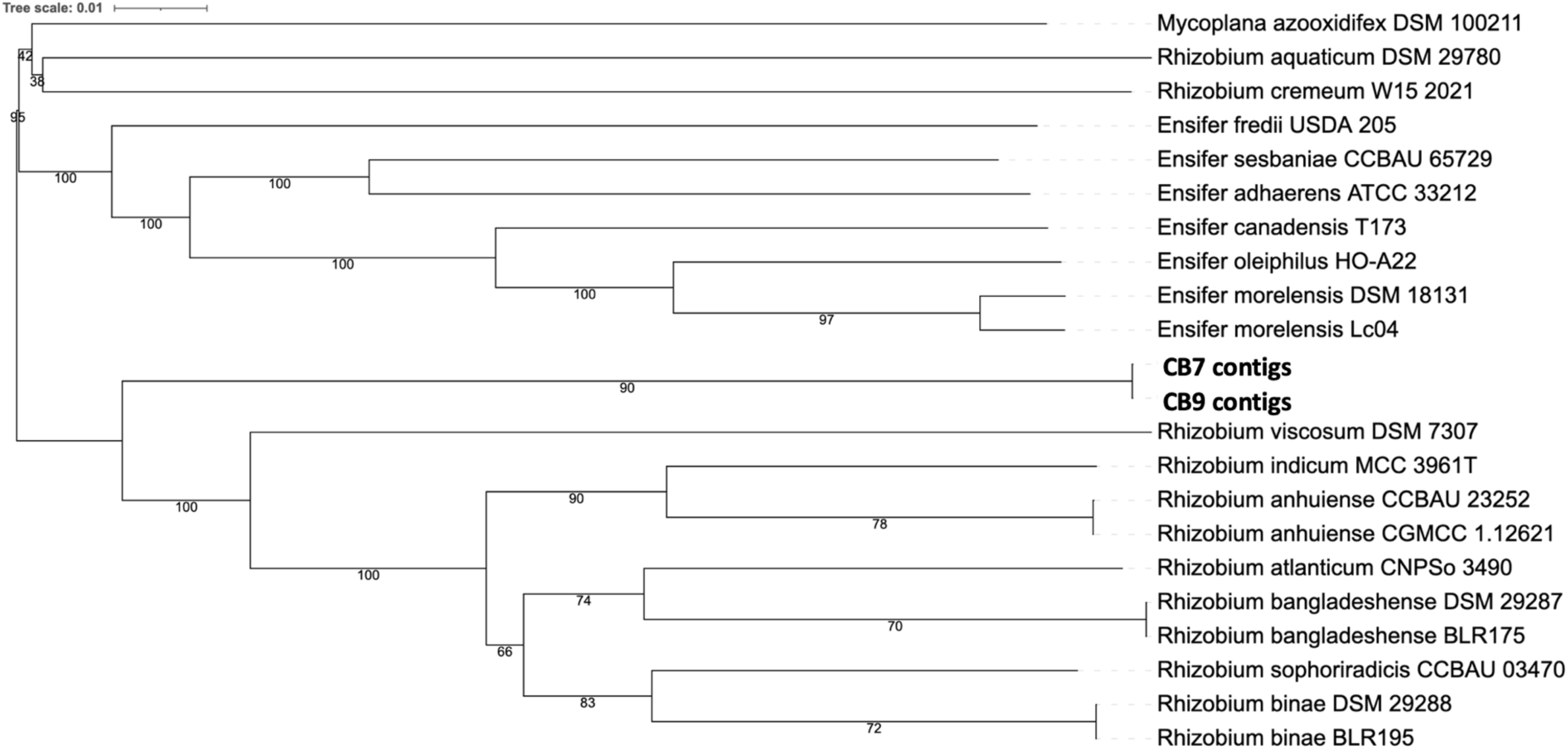
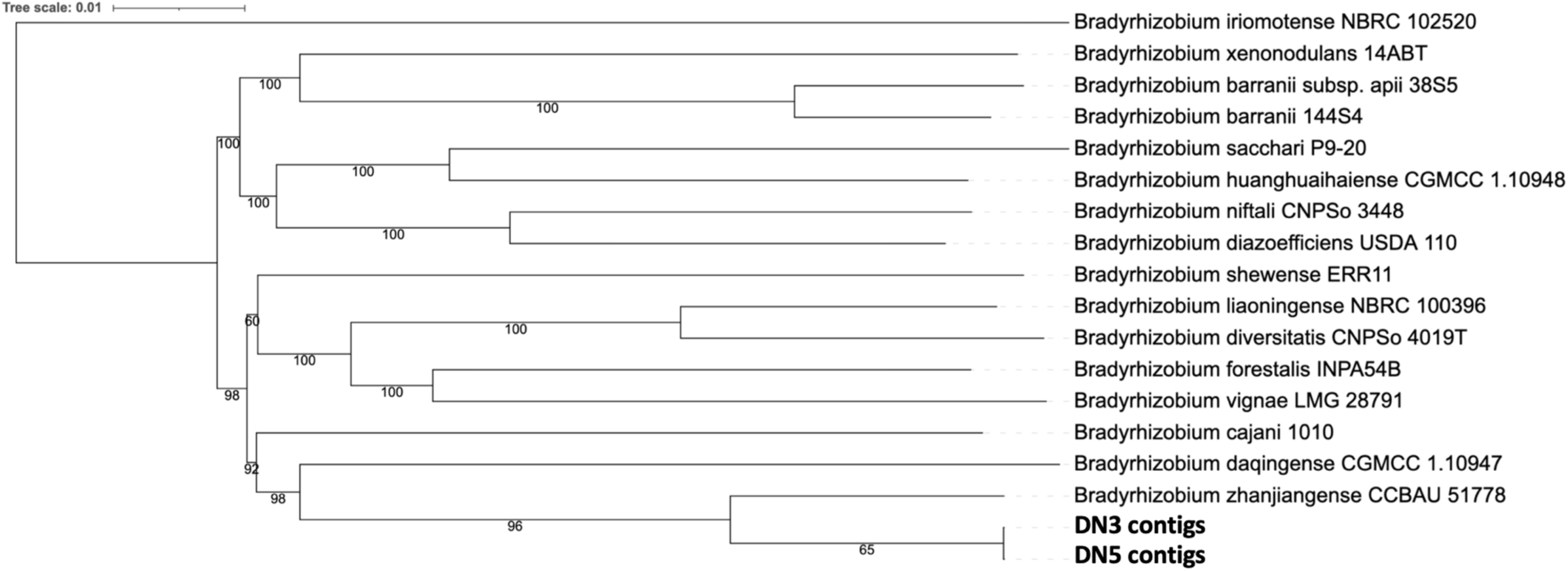
A: Tree showing relative distance of CB7 and CB9, which were isolated from *V. unguiculata*. **B:** Tree showing relative distance of DN3 and DN5, which were isolated from *L. purpureus* (48)(49).

**Figure 3:**
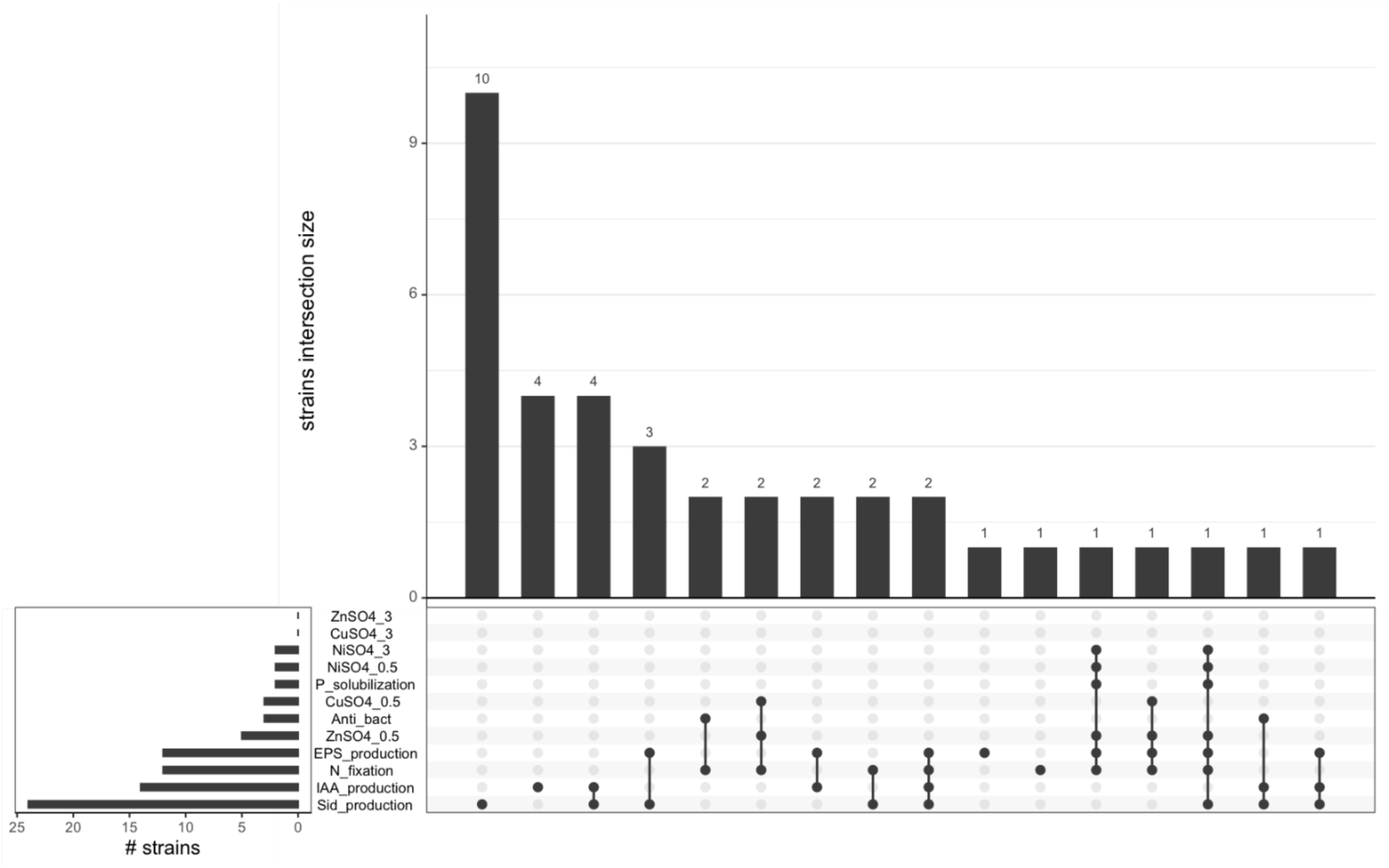
Diagrammatic presentation of the *in vitro* phenotypes of the root nodule bacterial endophytes strains. Each bar chart in the upset plot represents the number of isolates showing a single or a combination of phenotypes. Phenotypes are arranged from the most common (Siderophore production) to the less distributed (ability to grow on ZnSo_4_ at 3 mg/mL). The different phenotypes tested, related to PGP effects, are explained as follow: Sid_production (ability to synthetize siderophores); IAA_production (ability to synthetize indole-3-acetic acid); N_fixation (ability to fix atmospheric nitrogen); EPS_production (ability to produce exopolysaccharides); ZnSO_4_ 0.5 mg/mL or 3 mg/mL (ability to grow in media supplemented with ZnSO_4_ at two different concentrations); Anti_Bac (antimicrobial activity against E. coli as target; CuSO_4_ 0.5 mg/mL or 3 mg/mL (ability to grow in media supplemented with CuSO_4_ at two different concentrations); P_solubilization (ability to solubilize phosphate); NiSO_4_ 0.5 mg/mL or 3 mg/mL (ability to grow in media supplemented with NiSO_4_).

Concerning the symbiotic and nitrogen-fixing genes, the genes, *nodC* and *nifH*, which are required for legume nodulation and nitrogen fixation, were surprisingly absent from the genome annotations of *Rhizobium* sp. CB7 and CB9 strains (despite them inducing nodulation -Suplementary Table 4), while they were present in *Bradyrhizobium* sp. DN3 and DN5 genomes. We cannot exclude that that the lack of identification of *nodC* and *nifH* could be due to the draft assembly or to limits of the current annotation. An additional search for *nodC* and *nifH* on CB7 and CB9 strains by BLAST analysis with protein orthologs from *Rhizobium indicum* as the closest species, (QKK34725.1 chitooligosaccharide synthase *nodC* and QKK34534.1 nitrogenase iron protein *nifH*) resulted in a partial identification of the DFAST LOCUS_33740 (annotated as glycosyltransferase family 2 protein) sharing similarity with the *nodC* sequence. The level of sequence identity between *Rhizobium indicum nodC* and LOCUS_33740 in CB7 is 28.17%, with a query coverage of 41% of the *nodC* protein. These values are relatively low and suggest that LOCUS_33740 is not an ortholog, but rather a distant paralog within the glycosyltransferase family.

However, the gene locus was not clustered with the *nodA* gene, as expected. No hits with *nifH* were found (cutoff for e-value 0.001) and no hits for orthologs of *nodC* and *nifH* were found in PGAP annotation files from GenBank. Re-annotation with KAAS (Kegg Automatic Annotation Server) on 2024.04.18., again reported no hits. From the genome sequence data we obtained, we can reasonably conclude that CB7 and CB9 strains do not harbour nitrogenase genes and could not synthesise canonical *nod* factors.

### Secondary metabolite gene clusters

Given the potential of these rhizobial isolates as novel bioinoculants, we further explored their secondary metabolite gene clusters, which could contribute to plant growth promotion or microbial antagonistic activities beyond their symbiotic roles with legumes. Both CB7 and CB9 were found to harbour a gene cluster responsible for the biosynthesis of vicibactin, displaying 100% similarity to the cluster in *Rhizobium etli* CFN 42. Vicibactin, a high-affinity iron chelator, is also present in *R. etli* and *R. leguminosarum* (68). Additionally, DN5 and DN3 possessed a non-ribosomal peptide synthetase gene cluster identical to the rhizomide A, B, and C gene cluster of *Paraburkholderia rhizoxinica* HKI 454. Rhizomides are known depsipeptide macrolactones with antimicrobial properties (69).

### Detection of putative antibiotic resistance determinants

The CARD database was then used to analyse the presence of systems putatively involved in antibiotic resistance. Results are in the Dataset CARD. As expected, most of the retrieved systems belong to the RND (Resistance Nodulation Cell Division) family which can determine multi drug resistance thanks to the efflux mechanisms. DN3 and DN5 strains also harboured a putative beta lactamase gene, involved in carbapenem resistance.

### Culture-dependent isolation and identification of non-rhizobial nodule-associated bacteria

The non-rhizobial nodule-associated bacteria isolated and identified in this study belonged to three phyla, the Firmicutes (also known as Bacillota) (88%), Actinomycetota (also known as Actinobacteria) 7% and Proteobacteria (5%). The majority belonged to the phylum Firmicutes and were taxonomically assigned to *Bacillus*, *Priestia, Paenibacillus, Gottfriedia, Neobacillus, Lysinibacillus, Fictibacillus*, and *Brevibacillus* (Supplementary Table 1).

Interestingly, one of the bacterial isolates was *Variovorax paradoxus* which was isolated from *V. unguiculata*; to the best of our knowledge, this is the first report of this bacterial species isolated from legumes in Namibia. *Variovorax* belongs to the family Comamonadaceae, within the order Burkholderiales. The V3–V4 region of the 16S rRNA gene offers limited resolution, making it difficult to distinguish between closely related genera or species. As a result, sequences may be incorrectly assigned to the broader Burkholderia-group due to insufficient phylogenetic resolution. In contrast, the full-length 16S rRNA gene obtained from isolates allows for more accurate and reliable taxonomic classification, and this probably the reason why we were not able to resolve the phylogeny better (70).

### Evaluation of PGP activity of non-rhizobial bacterial nodule isolates

A set of plant-growth-promoting (PGP) associated phenotypes were performed *in vitro* on the non-rhizobial strains isolated in this study; these included nitrogen fixation, indole acetic acid (IAA) production, siderophore production, antifungal activity against *F. graminearum*, exopolysaccharide production, and phosphate solubilisation. Results are summarised in Supplementary Table 3 and also shown diagramatically in Figure 3.

Siderophore production was observed in 23 isolates belonging to all three phyla. This included *Bacillus, Paenibacillus*, and *Streptomyces* genera. Nitrogen fixation potential was observed in eight isolates from the genera *Priestia, Paenibacillus* and *Xanthomonas*. Fourteen isolates were observed to produce IAA; the highest concentration was observed from a *Brevibacillus* unclassified species isolated from *V. aconitifolia* IPCMO-880.

Only three strains displayed anti-fungal activity against *F. graminearum*; two *Paenibacillus* species (including *Paenibacillus polymyxa*) and a *Bacillus subtilis subsp. subtilis*. Ten isolates showed exopolysaccharide production mostly isolated from *V. unguiculata* and *V. radiata*. The EPS producing isolates were identified as *Priestia* species *P. aryabhattai, P. filamentosa* and two unclassified species. The remaining isolates were from the *Brevibacillus,* and *Xanthomonas* genera. To conclude, none of the isolates showed the ability to solubilize inorganic phosphates after 7 days. The *V. paradoxus* isolate reported in this study was positive for EPS production and nitrogen fixation but not for IAA production. Additionally, this strain was tolerant to 0.5 mg/mL ZnSO_4_ and 0.5mg/ml of CuSO_4_.

## Discussion

This work presented data on bacterial communities from root nodules of five legume species grown in Namibia and also constitute a culture collection of native rhizobial and non-rhizobial legume-associated bacteria with the long-term aims to use some of these strains as bioinoculants to increase productivity and resilience of legume crops in Namibia and other arid regions in Africa. The study of the biodiversity of soil bacteria which may associate with plants is now a major topic in the search for sustainable solutions to food and nutrient security and maintain agriculture productivity under climate change scenarios (22). Particularly, it is timely to study the effect of locally isolated bacteria as bioinoculants which possess beneficial traits that are well-suited to specific challenges or stressors present in a specific environment and that can maximize their compatibility with the local conditions and increase the potential for positive effects on plant growth and health.

Bacterial community studies revealed that the *Bradyrhizobium* genus was predominant in the nodules community, as expected. Previous studies have isolated *Bradyrhizobium* sp. in *V. subterraenea* (71), *V. unguiculata* (72)*, Glycine clandestine, G. max* (73) and *L. purpureus* (74). In particular, the two most abundant bacterial families identified in this study were the Nitrobacteraceae and Xanthobacteraceae; these are the families in which the rhizobia genera are found. Importantly, the genus *Bradyrhizobium* has been recently re-classified under the family nomenclature Bradyrhizobiaceae. In some cases, Bradyrhizobiaceae has been named as Nitrobacteraceae citing the more fitting nitrogen<u>-</u>fixing ability of the species in this family (75). The results obtained in this study were similar to a previous analysis of the nodule microbiota of soybean (*G. max*) since *Bradyrhizobium* species contributed more than 99% of the Proteobacteria sequences (76). On the other hand, the genus *Rhizobium*, was not detected in *V. unguiculata* in the microbiota studies despite being isolated from the nodules of this legume. This may be attributed to the parameters set for low abundant taxa (<50 reads per sample were excluded).

The relative abundance of the taxa taxonomically identified as rhizobia showed significant differences across the different legume species. This could be due to rhizobia being specific to host plants (77,78). Furthermore, the different microbial compositions which have been detected in nodules may be linked to the presence of both biotic and abiotic stress factors. These stress factors may affect nodulation, microbial colonisation, and composition (79). This was not factored in our study, as no plants showed physiological evidence of stress.

Other non-rhizobial families were identified from the nodule-extracted DNA. The most abundant and largely present family was Bacillaceae, which includes the *Bacillus* genus that is a commonly observed non-rhizobial nodule occupant (80). Studies have shown that root nodules harbour a microbiota which is composed not only by symbiotic nitrogen-fixing rhizobia, but by also of a multitude of other species which may behave as commensal endophytes which can also affect rhizobial growth and activity (10). Consequently, it was interesting to isolate non-rhizobial bacterial strains from the root nodules of several legume species grown in Namibia as described in the Materials and Methods section.

Several species belonging to *Bacillus* can promote plant health, regulate nodule formation (78) and exhibit antagonistic activity against plant pathogens (81), suggesting a possible role of strains belonging to this genus as having positive interactions with rhizobial strains. In addition, the Burkholderiaceae family has been observed in this study as another nodule occupant; this result is in line with a previous study documenting their presence in root nodules (82). These data agree with the collection of non-rhizobial isolates identified, which were largely belonging to the Bacillota (syn. Firmicutes) phylum (83) in the families *Bacillaceae* and *Paenibacillaceae* (84).

However, further analysis, employing also fluorescently tagged strains should be performed to confirm the tissue localization of such bacterial isolates, the role of their PGP properties and their interaction with the nodule development and rhizobial symbiotic nitrogen fixation.

The phylogenomic characterization of the four rhizobial strains isolated in this study revealed that they belonged to the genera *Rhizobium* (CB7 and CB9) and *Bradyrhizobium* (DN3 and DN5). Based on G + C content, the isolates CB7 and CB9 are likely to be either the same species or highly similar strains of the same species; a similar conclusion can be drawn for the isolates DN3 and DN5. Nevertheless, the dDDH values for both genera indicate that the strain genomes may be novel (previously undescribed) species (72,85). Genome mining surprisingly noted the absence of *nodC* and *nifH* genes from CB7 and CB9 genomes; this may be attributed to the draft assembly or to limits of the current annotation.

However, the *Rhizobium* strains (CB7 and CB9) induced significantly less nodules compared to the *Bradyrhizobium* strains. This could be attributed to the absence of *nodC* genes in the rhizobium strains (86). However, the observation of nodulation in the absence of *nodC* genes by CB7 and CB9 strains may suggest the possibility of alternative nodulation pathways, as already reported for photosynthetic bradyrhizobia (87). We cannot consequently *a priori* exclude that *Rhizobium* CB7 and CB9 strains may harbour some novel or atypical nodulation pathway. It is important to note that the observed nodules in *Rhizobium* inoculated plants may have been due to other *nod* genes as previously described (88). It is also important to note that the nodules were not assessed for their ability to fix nitrogen.

Concerning other bradyrhizobia isolated from Namibian soils, *B. vignae* was distant from the DN3 and DN5 strains. For the other bradyrhizobia isolated from Namibia (*B. kavangense*, *B. subterraneum*, *B. namibiense* (74,89), no genome sequences are currently available hampering a full genome comparison and deep taxonomic analysis (16S rRNA gene sequences are not very informative for species delineation in the genus *Bradyrhizobium* (89)). *Bradyrhizobium* sp. DN3 and *Bradyrhizobium* sp. DN5 had a genome size of ca. 8.4 Mbp, a GC% content of 63.2%, a single rRNA operon and ca. 7900 genes, similarly to other bradyrhizobia (90). *Rhizobium* sp. CB7 and *Rhizobium* sp. CB9 a genome size of ca. 7.6 Mbp, ca. 7900 genes, a single rRNA operon and a GC content% of 59.6%. Given the high genomic diversity of member of the *Rhizobium* genus, genome sizes of CB9 and CB7 lie within the upper values for member of this genus (91).

Further studies will also assess their beneficial effects and their potential as bioinoculants for biofertilizer development in field trials in Namibia and other drylands.

## Supporting information

https://drive.google.com/file/d/1nZ3kS_TbOj81cZ0kOMG0kJ0cP0H116l1/view?usp=drive_link

## Acknowledgments

The authors would like to express their gratitude for the support by the International Centre for Genetic Engineering and Biotechnology to Dr Vittorio Venturi’s Bacteriology Laboratory. We thank Ms Iris Bertani for her support in the planning and development of the experiment. Paidamoyo Mataranyika was also graciously supported by the Women Scientists in Africa (WE-STAR) fellowship under the Italian Ministry of Foreign Affairs and International Cooperation (MAECI) and the International Centre for Genetic Engineering and Biotechnology (ICGEB).

## Author Contributions

Conceptualization, P.N.M, P.M.C, V.V, and J.D.U; Methodology, P.N.M, O.S.O, V.V; Formal Analysis, P.N.M, AM, FV, CB; Investigation, P.N.M; Writing – Original Draft Preparation, P.N.M; Writing – Review & Editing, P.M.C, V.V, O.S.O, A.M, F.V, CB J.D.U.; Supervision, P.M.C, V.V, and J.D.U; Project Administration, V.V, P.M.C, and J.D.U.

## Ethical statement

*Six accessions from five legume species* were used in this study. These are dolichos [*Lablab purpureus* (L.) Sweet var. Lignosus Prain] accession IC0623025, mung bean [Vigna radiata (L.) R. Wilczek var. radiata] accession IC39399, cowpea (Vigna unguiculata L. Walp) accession Gujarat 5, horsegram (*Macrotyloma uniflorum* Var. Madhu) accession Himala, and mothbean [*Vigna aconitifolia* (Jacq.) Marechal] accessions RMB-25 and IPCMO-880. They were obtained from the National Bureau of Plant Genetic Resources’ (NBPGR), India in July 2019.

## Data availability

Genome sequences presented in this study are available under BioProject PRJNA896769. Accession numbers and additional data are presented in Supplementary material.

## Supplementary data

Data is contained within the article or supplementary material.

---

**Supplementary Table 1:** Assigned accession numbers for the partial 16S rRNA gene sequence identities.

| Sample ID | Assigned sequence numbers |
| --- | --- |
| **Rhizobial strains** |  |
| CB7 | OR619527 |
| CB9 | OR619528 |
| DN3 | OR619529 |
| DN5 | OR619530 |
| **Non-rhizobial strains** |  |
| CN5 | OR619526 |
| C11 | OP623457 |
| C3 | OP623475 |
| C1 | OP623474 |
| C5 | OP623456 |
| C6 | OP623476 |
| CA9 | OP623476 |
| D10 | OP623458 |
| D14 | OP623479 |
| D7 | OP623488 |
| D5 | OP623476 |
| D17.1 | OP623480 |
| MB17 | OP623491 |
| MB16 | OP623473 |
| MB1 | OP623469 |
| MB14 | OP623472 |
| MB11 | OP623471 |
| MB8 | OP623470 |
| H14 | OP623481 |
| H13 | OP623460 |
| H4 | OP623459 |
| M25-17 | OP623490 |
| M25-11 | OP623492 |
| M25-13 | OP623467 |
| M25-12 | OP623466 |
| M25-10 | OP623493 |
| M25-16 | OP623468 |
| M25-6 | OP623489 |
| M25-5 | OP623465 |
| M25-7 | OP623494 |
| M25-3 | OP623464 |
| M8-2 | OP623461 |
| M8-8 | OP623482 |
| M8-6 | OP623462 |
| M8-21 | OP623463 |
| M8-14 | OP623485 |
| M8-9 | OP623483 |
| M8-13 | OP623484 |
| M8-16.1 | OP623486 |
| M8-16.2 | OP623487 |

**Supplementary Table 2:** Assigned accession numbers for sequence reads.

| Sample ID | Legume source | Assigned Accession |
| --- | --- | --- |
| D1 | *Lablab purpureus* | SAMN31566282 |
| D2 | *Lablab purpureus* | SAMN31566283 |
| D3 | *Lablab purpureus* | SAMN31566284 |
| C1 | *Vigna unguiculata* | SAMN31566285 |
| C2 | *Vigna unguiculata* | SAMN31566286 |
| C3 | *Vigna unguiculata* | SAMN31566287 |
| H1 | *Macrotyloma uniflorum* | SAMN31566288 |
| H2 | *Macrotyloma uniflorum* | SAMN31566289 |
| H3 | *Macrotyloma uniflorum* | SAMN31566290 |
| M8-2 | *Vigna aconitifolia* | SAMN31566291 |
| M8-3 | *Vigna aconitifolia* | SAMN31566292 |

**Supplementary Table 3:** Plant growth-promoting traits of root nodule isolates.

| Legume | Common name | Accession | Sample | Bacterial genus/species | Phos<sup>a</sup> | Sid<sup>b</sup> | Nit<sup>c</sup> | Ant<sup>d</sup> | EPS<sup>e</sup> | IAA<sup>f</sup> |
| --- | --- | --- | --- | --- | --- | --- | --- | --- | --- | --- |
| *Lablab purpureus* | Dolichos | IC0623025 | D10 | *Priestia sp.* | - | - | - | - | - | - |
|  |  |  | D14 | *Bacillus sp.* | - | + | - | - | - | - |
|  |  |  | D7 | *Paenibacillus sp.* | - | - | - | - | - | - |
|  |  |  | D5 | *Priestia sp.* | - | + | + | - | - | - |
|  |  |  | D17.1 | *Bacillus subtilis subsp. subtilis* | un | + | un | + | - | + |
| *Vigna unguiculata* | Cowpea | Gujarat 5 | C1 | *Paenibacillus sp.* | - | + | - | - | + | - |
|  |  |  | C3 | *Priestia sp.* | - | - | - | - | + | + |
|  |  |  | C11 | *Priestia aryabhattai* | - | - | - | - | + | - |
|  |  |  | C5 | *Gottfriedia luciferensis* | - | + | - | - | + | + |
|  |  |  | C6 | *Priestia sp.* | - | - | - | - | - | - |
|  |  |  | CA9 | *Streptomyces sp.* | - | + | - | - | - | - |
|  |  |  | CA4 | *Streptomyces sp.* | - |  | - | - | - | - |
| *Macrotyloma uniflorum* | Horsegram | Himala | H14 | *Priestia aryabhattai* | - | - | - | - | Un | + |
|  |  |  | H13 | *Neobacillus sp.* | - | + | - | - | - | - |
|  |  |  | H4 | *Priestia sp.* | - | - | - | - | Un | + |
| *Vigna radiata* | Mungbean | IC39399 | MB17 | *Priestia aryabhattai* | - | - | - | - | - | - |
|  |  |  | MB16 | *Priestia filamentosa* | - | + | - | - | + | - |
|  |  |  | MB1 | *Xanthomonas sp.* | - | + | + | - | + | + |
|  |  |  | MB14 | *Paenibacillus sp.* | - | - | + | + | - | - |
|  |  |  | MB11 | *Bacillus sp.* | - | + | - | - | - | - |
|  |  |  | MB3.1 | *Priestia sp.* | - | + | + | - | + | + |
|  |  |  | MB8 | *Gottfriedia sp.* | - | + | - | - | + | - |
| *Vigna aconitifolia* | Moth bean | RMB-25 | M25-17 | *Gottfriedia luciferensis* | - | - | - | - | - | - |
|  |  |  | M25-11 | *Lysinibacillus boronitolerans* | - | - | - | - | - | + |
|  |  |  | M25-13 | *Paenibacillus sp.* | - | + | - | - | - | - |
|  |  |  | M25-12 | *Xanthomonas sp.* | - | + | - | - | - | - |
|  |  |  | M25-10 | *Priestia aryabhatai* | - | - | + | - | - | - |
|  |  |  | M25-16 | *Gottfriedia luciferensis* | - | - | - | - | - | - |
|  |  |  | M25-6 | *Xanthomonas campestris* | - | + | + | - | - | - |
|  |  |  | M25-5 | *Fictibacillus sp.* | - | + | - | - | - | - |
|  |  |  | M25-7 | *Fictibacillus sp.* | - | + | - | - | - | - |
|  |  |  | M25-3 | *Priestia aryabhatai* | - | - | - | - | - | - |
|  | Moth bean | IPCMO-880 | M8-2 | *Priestia sp.* | - | + | - | - | - | - |
|  |  |  | M8-8 | *Neobacillus sp.* | - | + | - | - | - | + |
|  |  |  | M8-6 | *Neobacillus sp.* | - | + | - | - | - | - |
|  |  |  | M8-21 | *Bacillus sp.* | - | + | - | - | - | + |
|  |  |  | M8-14 | *Paenibacillus polymyxa* | - | - | + | + | - | - |
|  |  |  | M8-9 | *Priestia aryabhatai* | - | - | - | - | - | + |
|  |  |  | M8-13 | *Brevibacillus sp.* | - | - | - | - | + | + |
|  |  |  | M8-16.1 | *Bacillus sp.* | - | + | - | - | - | + |
|  |  |  | M8-16.2 | *Bacillus proteolyticus* | - | + | - | - | - | + |

<sup>a</sup> Phosphate solubilization
<sup>b</sup> Siderophore production
<sup>c</sup> Nitrogen fixation
<sup>d</sup> Antifungal activity
<sup>e</sup> Exopolysaccharide production
<sup>f</sup> Indole acetic acid production
“-” means showed no production/ “+” means showed production/ “un” means plant growth-promoting trait could not be determined.

**Supplementary Table 4:** Effect of Rhizobia on nodulation.

| **Control: No inoculant** |  |  |  | **DN3: *Bradyrhizobium*** |  |  |  |
| --- | --- | --- | --- | --- | --- | --- | --- |
| Number of Germinated plants out of 12 |  | 9 |  | Number of Germinated plants out of 12 |  | 8 |  |
| Number of plants with nodules |  | 0 |  | Number of plants with nodules |  | 8 |  |
| Plant | # of pods/plant | # of seeds/pod | # of nodules/plant | Plant | # of pods/plant | # of seeds/pod | # of nodules/plant |
| 1 | 0 | 0 | 0 | 1 | 1 | 5 | >10 |
| 2 | 0 | 0 | 0 | 2 | 1 | 4 | >10 |
| 3 | 1 | 5 | 0 | 3 | 2 | 3 | >10 |
| 4 | 1 | 3 | 0 | 4 | 1 | 2 | >10 |
| 5 | 2 | 2 | 0 | 5 | 1 | 5 | >10 |
| 6 | 1 | 2 | 0 | 6 | 1 | 4 | >10 |
| 7 | 2 | 2 | 0 | 7 | 1 | 2 | >10 |
| 8 | 1 | 2 | 0 | 8 | 2 | 3 | >10 |
| 9 | 1 | 5 | 0 | 9 |  |  |  |
| 10 |  |  |  | 10 |  |  |  |
| 11 |  |  |  | 11 |  |  |  |
| 12 |  |  |  | 12 |  |  |  |
| **CB7 *Rhizobium*** |  |  |  | **DN5 *Bradyrhizobium*** |  |  |  |
| Number of Germinated plants out of 12 |  | 12 |  | Number of Germinated plants out of 12 |  | 7 |  |
| Number of plants with nodules |  | 3 |  | Number of plants with nodules |  | 7 |  |
| Plant | # of pods/plant | # of seeds/pod | # of nodules/plant | Plant | # of pods/plant | # of seeds/pod | # of nodules/plant |
| 1 | 1 | 9 | 0 | 1 | 0 | 0 | >10 |
| 2 | 1 | 7 | 0 | 2 | 0 | 0 | >10 |
| 3 | 1 | 3 | 0 | 3 | 0 | 0 | >10 |
| 4 | 1 | 4 | 4 | 4 | 0 | 0 | >10 |
| 5 | 1 | 2 | 0 | 5 | 0 | 0 | >10 |
| 6 | 1 | 2 | 0 | 6 | 2 | Juvenile | >10 |
| 7 | 1 | 4 | 0 | 7 | 1 | Juvenile | >10 |
| 8 | 1 | 3 | 0 | 8 |  |  |  |
| 9 | 0 | 0 | 1 | 9 |  |  |  |
| 10 | 1 | 2 | 0 | 10 |  |  |  |
| 11 | 1 | Juvenile | 0 | 11 |  |  |  |
| 12 | 0 | 0 | 2 | 12 |  |  |  |
| **CB9 *Rhizobium*** |  |  |  | **CN5 *V. paradoxus*** |  |  |  |
| Number of Germinated plants out of 12 |  | 6 |  | Number of Germinated plants out of 12 |  | 11 |  |

| Number of plants with nodules |  |  | 3 | Number of plants with nodules |  |  | 11 |
| --- | --- | --- | --- | --- | --- | --- | --- |
| Plant | # of pods/plant | # of seeds/pod | # of nodules/plant | Plant | # of pods/plant | # of seeds/pod | # of nodules/plant |
| 1 | 1 | 4 | 0 | 1 | 1 | Juvenile | >10 |
| 2 | 1 | 5 | 5 | 2 | 1 | Juvenile | >10 |
| 3 | 1 | Juvenile | 0 | 3 | 1 | 4 | >10 |
| 4 | 0 | 0 | 0 | 4 | 0 | 0 | >10 |
| 5 | 1 | 4 | 1 | 5 | 1 | Juvenile | >10 |
| 6 | 1 | 4 | 7 | 6 | 2 | Juvenile | >10 |
| 7 |  |  |  | 7 | 0 | 0 | >10 |
| 8 |  |  |  | 8 | 1 | Juvenile | >10 |
| 9 |  |  |  | 9 | 1 | Juvenile | >10 |
| 10 |  |  |  | 10 | 0 | 0 | >10 |
| 11 |  |  |  | 11 | 2 | Juvenile | >10 |
| 12 |  |  |  | 12 |  |  |  |

**Supplementary Table 5.** Results of ANIb%. In brackets the % of aligned nucleotides.

|  | CB7 | CB9 | DN3 | DN5 |
| --- | --- | --- | --- | --- |
| CB7 | * | 99.99 [99.83] | 67.37 [18.96] | 67.36 [18.97] |
| CB9 | 99.98 [99.85] | * | 67.54 [18.97] | 67.53 [18.98] |
| DN3 | 67.35 [16.89] | 67.35 [16.89] | * | 99.99 [99.64] |
| DN5 | 67.38 [16.97] | 67.36 [16.97] | 99.99 [99.63] | * |

