## Supplementary material for "Bacterial community studies and novel *Bradyrhizobium* and *Rhizobium* strains from drought-tolerant legumes grown in Namibia": https://drive.google.com/file/d/1nZ3kS_TbOj81cZ0kOMG0kJ0cP0H116l1/view?usp=drive_link

Ty

| TYGS ID | Kind | Species clu | Subspecies | Preferred name | Deposit | Authority | Other deposits |
|---|---|---|---|---|---|---|---|
| 121491 | type strain | 1 | 0 | Pararhizobium capsulatu | DSM 1112 | (Hirsch and Müller 1986) Mo | ATCC 43294; DSM 112; V |
| 126532 | type strain | 2 | 4 | Ensifer morelensis | DSM 18131 | (Wang et al. 2002) Wang et a | LMG 21331; DSM 18131; |
| 156842 | type strain | 2 | 4 | Ensifer morelensis | Lc04 | (Wang et al. 2002) Wang et a | LMG 21331; DSM 18131; |
| 13653 | type strain | 3 | 1 | Rhizobium viscosum | DSM 7307 | (Gasdorf et al. 1965) Flores-F | LMG 16473; CIP 82.105; |
| 143792 | type strain | 4 | 2 | Rhizobium aquaticum | DSM 29780 | Máthé et al. 2019 | JCM 31760; SA-276 |
| 147570 | type strain | 5 | 3 | Rhizobium cremeum | W15(2021) | Yang et al. 2022 | CGMCC 1.18731; KACC 2 |
| 158005 | type strain | 6 | 5 | Rhizobium chutanense | C5T | Huo et al. 2019 | LMG 30777; CCTCC AB 2 |
| 19935 | type strain | 7 | 6 | Ensifer sesbaniae | CCBAU 65729 | Wang et al. 2015 | LMG 26833; HAMBI 3287 |
| 20793 | type strain | 8 | 7 | Rhizobium sophoriradici | CCBAU 03470 | Jiao et al. 2015 | LMG 27898; C-5-1; HAM |
| 3420 | type strain | 9 | 8 | Ensifer fredii | USDA 205 | (Scholla and Elkan 1984) You | LMG 6217; NRRL B-1424 |
| 3743 | type strain | 10 | 9 | Ensifer adhaerens | ATCC 33212 | Casida 1982 emend. Hördt e | A; LMG 20216; JCM 2110 |
| 13215 | type strain | 11 | 10 | Rhizobium anhuiense | CCBAU 23252 | Zhang et al. 2015 | LMG 27729; CGMCC 1.12 |
| 48401 | type strain | 11 | 10 | Rhizobium anhuiense | CGMCC 1.1262 | Zhang et al. 2015 | LMG 27729; CGMCC 1.12 |
| 52434 | type strain | 12 | 11 | 'Rhizobium indicum' | MCC 3961T | Rahi et al. 2020 | JCM 33658; JKLM 12A2; |
| 7640 | type strain | 13 | 12 | Mycoplana azooxidifex | DSM 100211 | (Behrendt et al. 2016) Kuzma | LMG 28788; Po 20/26 |
| 143781 | type strain | 14 | 13 | Rhizobium bangladeshe | DSM 29287 | Rashid et al. 2015 | LMG 28442; DSM 29287; |
| 81029 | type strain | 14 | 13 | Rhizobium bangladeshe | BLR175 | Rashid et al. 2015 | LMG 28442; DSM 29287; |
| 126819 | type strain | 15 | 14 | Rhizobium binae | DSM 29288 | Rashid et al. 2015 | LMG 28443; DSM 29288; |
| 81033 | type strain | 15 | 14 | Rhizobium binae | BLR195 | Rashid et al. 2015 | LMG 28443; DSM 29288; |
| 94614 | type strain | 16 | 15 | Ensifer canadensis | T173 | Bromfield et al. 2023 | LMG 32374; HAMBI 3766 |
| U519341 | user strain | 17 | 16 | CN7 contigs | | | |

ype Strain Genome Server

| Synonymous taxon names | Base pairs | Percent G+C | No. protei | Goldstamp | Bioproject acces | Biosample access | Assembly accessi | IMG OID | BacDive |
|---|---|---|---|---|---|---|---|---|---|
| Blastobacter capsulatus; Para | 6477857 | 5934 | 6290 | Gp0251907 | | | | 2927149029 | strain page |
| Ensifer morelensis; Sinorhizob | 6824949 | 6186 | 6407 | Gp0505705 | | | | 2913284250 | strain page |
| Ensifer morelensis; Sinorhizob | 7066229 | 6175 | 6549 | | PRJNA224116 | SAMN14518349 | GCF_013283195 | | strain page |
| Arthrobacter viscosus; Rhizob | 7069622 | 6001 | 6898 | Gp0456036 | | | | 2870861536 | strain page |
| Rhizobium aquaticum | 4903511 | 61 | 4687 | Gp0538768 | | | | 2928412209 | strain page |
| Rhizobium cremeum | 5328908 | 6166 | 5113 | | PRJNA224116 | SAMN18012446 | GCF_022884065 | | species page |
| Rhizobium chutanense | 6873329 | 6139 | 6474 | | PRJNA224116 | SAMN07627103 | GCF_002531935 | | species page |
| Ensifer sesbaniae | 6895946 | 621 | 6457 | | PRJNA622509 | SAMN14518350 | GCA_013283665 | | species page |
| Rhizobium sophoriradicis | 6664732 | 6129 | 6158 | | PRJNA504373 | SAMN10390643 | GCA_003939025 | | species page |
| Ensifer fredii; Rhizobium fred | 6579820 | 6232 | 5932 | Gp0048750 | PRJNA211973 | SAMN04301590 | GCA_001461695 | | strain page |
| Ensifer adhaerens; Sinorhizob | 7280736 | 6233 | 6607 | Gp0094824 | PRJNA247817 | SAMN02781308 | GCA_000697965 | 2582581023 | strain page |
| Rhizobium anhuiense | 7116000 | 6113 | 6617 | | PRJNA503677 | SAMN10370526 | GCA_003985145 | | species page |
| Rhizobium anhuiense | 7110473 | 6111 | 6701 | | PRJDB10509 | SAMD00244951 | GCA_014638185 | | species page |
| Rhizobium indicum | 7533049 | 6078 | 7212 | | PRJNA224116 | SAMN11792776 | GCF_005862305 | | species page |
| Mycoplana azooxidifex; Rhizo | 5882088 | 6427 | 5545 | Gp0401155 | | | | 2829954201 | strain page |
| Rhizobium bangladeshense | 6302736 | 6093 | 6101 | Gp0538759 | | | | 2928359186 | strain page |
| Rhizobium bangladeshense | 6348611 | 6091 | 6047 | | PRJNA224116 | SAMN18219174 | GCF_017357245 | | strain page |
| Rhizobium binae | 6947669 | 611 | 6815 | Gp0538760 | | | | 2928991799 | strain page |
| Rhizobium binae | 7055288 | 6107 | 6716 | | PRJNA224116 | SAMN18219163 | GCF_017357225 | | strain page |
| Ensifer canadensis | 8094229 | 6096 | 7271 | | PRJNA713338 | SAMN18249011 | GCA_017488845 | | species page |
| | 7638399 | 5956 | 7292 | | | | | | |

Type Strain Genome Ser

| TYGS ID | Kind | Species clu | Subspecies | Preferred name | Deposit | Authority | Other deposits | Synonymous taxon n |
|---|---|---|---|---|---|---|---|---|
| 121491 | type strain | 1 | 0 | Pararhizobium capsulatu | DSM 1112 | (Hirsch and Müller 1 | ATCC 43294; DSM 112; VK | Blastobacter capsulat |
| 126532 | type strain | 2 | 4 | Ensifer morelensis | DSM 18131 | (Wang et al. 2002) V | LMG 21331; DSM 18131; N | Ensifer morelensis; Si |
| 156842 | type strain | 2 | 4 | Ensifer morelensis | Lc04 | (Wang et al. 2002) V | LMG 21331; DSM 18131; N | Ensifer morelensis; Si |
| 13653 | type strain | 3 | 1 | Rhizobium viscosum | DSM 7307 | (Gasdorf et al. 1965 | LMG 16473; CIP 82.105; N | Arthrobacter viscosus |
| 143792 | type strain | 4 | 2 | Rhizobium aquaticum | DSM 29780 | Máthé et al. 2019 | JCM 31760; SA-276 | Rhizobium aquaticum |
| 147570 | type strain | 5 | 3 | Rhizobium cremeum | W15(2021) | Yang et al. 2022 | CGMCC 1.18731; KACC 22 | Rhizobium cremeum |
| 158005 | type strain | 6 | 5 | Rhizobium chutanense | C5T | Huo et al. 2019 | LMG 30777; CCTCC AB 201 | Rhizobium chutanens |
| 19935 | type strain | 7 | 6 | Ensifer sesbaniae | CCBAU 65729 | Wang et al. 2015 | LMG 26833; HAMBI 3287 | Ensifer sesbaniae |
| 20793 | type strain | 8 | 7 | Rhizobium sophoriradici | CCBAU 03470 | Jiao et al. 2015 | LMG 27898; C-5-1; HAMBI | Rhizobium sophorirad |
| 3420 | type strain | 9 | 8 | Ensifer fredii | USDA 205 | (Scholla and Elkan 1 | LMG 6217; NRRL B-14241; | Ensifer fredii; Rhizobi |
| 3743 | type strain | 10 | 9 | Ensifer adhaerens | ATCC 33212 | Casida 1982 emend | A; LMG 20216; JCM 21105 | Ensifer adhaerens; Sir |
| 13215 | type strain | 11 | 10 | Rhizobium anhuiense | CCBAU 23252 | Zhang et al. 2015 | LMG 27729; CGMCC 1.126 | Rhizobium anhuiense |
| 48401 | type strain | 11 | 10 | Rhizobium anhuiense | CGMCC 1.12621 | Zhang et al. 2015 | LMG 27729; CGMCC 1.126 | Rhizobium anhuiense |
| 52434 | type strain | 12 | 11 | 'Rhizobium indicum' | MCC 3961T | Rahi et al. 2020 | JCM 33658; JKLM 12A2; K | Rhizobium indicum |
| 7640 | type strain | 13 | 12 | Mycoplana azooxidifex | DSM 100211 | (Behrendt et al. 201 | LMG 28788; Po 20/26 | Mycoplana azooxidife |
| 143781 | type strain | 14 | 13 | Rhizobium bangladeshen | DSM 29287 | Rashid et al. 2015 | LMG 28442; DSM 29287; E | Rhizobium banglades |
| 81029 | type strain | 14 | 13 | Rhizobium bangladeshen | BLR175 | Rashid et al. 2015 | LMG 28442; DSM 29287; E | Rhizobium banglades |
| 126819 | type strain | 15 | 14 | Rhizobium binae | DSM 29288 | Rashid et al. 2015 | LMG 28443; DSM 29288; E | Rhizobium binae |
| 81033 | type strain | 15 | 14 | Rhizobium binae | BLR195 | Rashid et al. 2015 | LMG 28443; DSM 29288; E | Rhizobium binae |
| 94614 | type strain | 16 | 15 | Ensifer canadensis | T173 | Bromfield et al. 202 | LMG 32374; HAMBI 3766 | Ensifer canadensis |
| U523333 | user strain | 17 | 16 | CN9 contigs | | | | |

ver

| Base pairs | Percent G- | No. protei | Goldstamp | Bioproject accessio | Biosample accessi | Assembly accessi | IMG OID | BacDive |
|---|---|---|---|---|---|---|---|---|
| 6477857 | 5934 | 6290 | Gp0251907 | | | | 2927149029 | strain page |
| 6824949 | 6186 | 6407 | Gp0505705 | | | | 2913284250 | strain page |
| 7066229 | 6175 | 6549 | | PRJNA224116 | SAMN14518349 | GCF_013283195 | | strain page |
| 7069622 | 6001 | 6898 | Gp0456036 | | | | 2870861536 | strain page |
| 4903511 | 61 | 4687 | Gp0538768 | | | | 2928412209 | strain page |
| 5328908 | 6166 | 5113 | | PRJNA224116 | SAMN18012446 | GCF_022884065 | | species page |
| 6873329 | 6139 | 6474 | | PRJNA224116 | SAMN07627103 | GCF_002531935 | | species page |
| 6895946 | 621 | 6457 | | PRJNA622509 | SAMN14518350 | GCA_013283665 | | species page |
| 6664732 | 6129 | 6158 | | PRJNA504373 | SAMN10390643 | GCA_003939025 | | species page |
| 6579820 | 6232 | 5932 | Gp0048750 | PRJNA211973 | SAMN04301590 | GCA_001461695 | | strain page |
| 7280736 | 6233 | 6607 | Gp0094824 | PRJNA247817 | SAMN02781308 | GCA_000697965 | 2582581023 | strain page |
| 7116000 | 6113 | 6617 | | PRJNA503677 | SAMN10370526 | GCA_003985145 | | species page |
| 7110473 | 6111 | 6701 | | PRJDB10509 | SAMD00244951 | GCA_014638185 | | species page |
| 7533049 | 6078 | 7212 | | PRJNA224116 | SAMN11792776 | GCF_005862305 | | species page |
| 5882088 | 6427 | 5545 | Gp0401155 | | | | 2829954201 | strain page |
| 6302736 | 6093 | 6101 | Gp0538759 | | | | 2928359186 | strain page |
| 6348611 | 6091 | 6047 | | PRJNA224116 | SAMN18219174 | GCF_017357245 | | strain page |
| 6947669 | 611 | 6815 | Gp0538760 | | | | 2928991799 | strain page |
| 7055288 | 6107 | 6716 | | PRJNA224116 | SAMN18219163 | GCF_017357225 | | strain page |
| 8094229 | 6096 | 7271 | | PRJNA713338 | SAMN18249011 | GCA_017488845 | | species page |
| 7645669 | 5956 | 7326 | | | | | | |

| TYGS ID | Kind | Species cluster | Subspecies cluster | Preferred name | Deposit | Authority |
|---|---|---|---|---|---|---|
| 104516 | type strain | 1 | 0 | 'Bradyrhizobium xenonodulans' | 14ABT | Claassens et al. 2023 |
| 115474 | type strain | 2 | 1 | Bradyrhizobium liaoningense | NBRC 100396 | Xu et al. 1995 |
| 14517 | type strain | 3 | 3 | 'Bradyrhizobium sacchari' | P9-20 | de Matos et al. 2017 |
| 15620 | type strain | 4 | 4 | 'Bradyrhizobium forestalis' | INPA54B | Martins da Costa et al. 2018 |
| 16415 | type strain | 5 | 5 | Bradyrhizobium daqingense | CGMCC 1.10947 | Wang et al. 2013 |
| 17297 | type strain | 6 | 6 | Bradyrhizobium cajani | 1010 | Araújo et al. 2017 |
| 18506 | type strain | 7 | 7 | Bradyrhizobium huanghuaihaiense | CGMCC 1.10948 | Zhang et al. 2012 |
| 18779 | type strain | 8 | 8 | Bradyrhizobium niftali | CNPSo 3448 | Klepa et al. 2019 |
| 18997 | type strain | 9 | 9 | Bradyrhizobium shewense | ERR11 | Aserse et al. 2018 |
| 2697 | type strain | 10 | 10 | Bradyrhizobium diazoefficiens | USDA 110 | Delamuta et al. 2013 |
| 120074 | type strain | 11 | 2 | Bradyrhizobium barranii subsp. apii | 38S5 | Bromfield et al. 2022 |
| 71334 | type strain | 11 | 11 | Bradyrhizobium barranii | 144S4 | Bromfield et al. 2022 |
| 8842 | type strain | 12 | 12 | 'Bradyrhizobium zhanjiangense' | CCBAU 51778 | Li et al. 2019 |
| 92658 | type strain | 13 | 13 | Bradyrhizobium diversitatis | CNPSo 4019T | Klepa et al. 2021 |
| 9288 | type strain | 14 | 14 | Bradyrhizobium vignae | LMG 28791 | Grönemeyer et al. 2016 |
| 94558 | type strain | 15 | 15 | Bradyrhizobium iriomotense | NBRC 102520 | Islam et al. 2010 |
| U523332 | user strain | 16 | 16 | DB3 contigs | | |

Type Strain Genome Server

| Other deposits | Synonymous taxon names | Base pairs | Percent G+C |
|---|---|---|---|
| LMG 31415; SARCC 753 | Bradyrhizobium xenonodulans | 8276659 | 6399 |
| 2281; LMG 18230; CIP 104858; ATCC 700350; DSM 24092 | Bradyrhizobium liaoningense | 7830169 | 6395 |
| BR 10280; HAMBI 3667 | Bradyrhizobium sacchari | 8685818 | 6384 |
| LMG 10044 | Bradyrhizobium forestalis | 8235644 | 6387 |
| LMG 26137; CCBAU 15774; HAMBI 3184 | Bradyrhizobium daqingense | 7885317 | 6374 |
| AMBPC1010; LMG 29967; CECT 9227 | Bradyrhizobium cajani | 8372827 | 6396 |
| LMG 26136; CCBAU 23303; HAMBI 3180 | Bradyrhizobium huanghuaihaiense | 9223029 | 6394 |
| CL 40; U687; USDA 10051 | Bradyrhizobium niftali | 9786055 | 6353 |
| LMG 30162; HAMBI 3532 | Bradyrhizobium shewense | 9162066 | 6323 |
| 3I1B110; ACCC 15034; CNPSo 46; BCRC 13528; CCRC 13528; NRR | Bradyrhizobium diazoefficiens | 9105828 | 6406 |
| LMG 31556; HAMBI 3721 | Bradyrhizobium barranii subsp. apii | 10530141 | 6334 |
| LMG 31552; HAMBI 3722 | Bradyrhizobium barranii; Bradyrhizobium barranii subsp. barranii | 11399526 | 6313 |
| LMG 29279; HAMBI 3648 | Bradyrhizobium zhanjiangense | 9342022 | 6294 |
| LMG 31650; WSM 4799 | Bradyrhizobium diversitatis | 8437014 | 6393 |
| 7-2; DSM 100297; NTCCM0018(Windhoek); NTCCM 18 | Bradyrhizobium vignae | 8169430 | 6322 |
| LMG 24129; EK05 | Bradyrhizobium iriomotense | 9736893 | 629 |
| | | 8458680 | 6319 |

| No. proteins | Goldstamp | Bioproject accession | Biosample accession | Assembly accession | IMG OID | BacDive |
|---|---|---|---|---|---|---|
| 7625 | | PRJNA680005 | SAMN16869375 | GCA_027594865 | | species page |
| 7615 | | PRJDB15044 | SAMD00576800 | GCA_030160735 | | strain page |
| 8163 | | PRJNA224116 | SAMN04849207 | GCF_002068095 | | species page |
| 7360 | | PRJNA418781 | SAMN08035463 | GCA_002795245 | | species page |
| 7442 | Gp0093889 | PRJNA255603 | SAMN02927912 | GCA_007830205 | | strain page |
| 7348 | | PRJNA593773 | SAMN13489115 | GCA_009759665 | | strain page |
| 8622 | Gp0093890 | PRJNA255602 | SAMN02927902 | GCA_007830635 | | strain page |
| 8664 | | PRJNA529074 | SAMN11254745 | GCA_004571025 | | species page |
| 8018 | Gp0108279 | PRJNA224116 | SAMN04487809 | GCF_900094605 | | species page |
| 8317 | Gp0000660 | PRJNA17 | SAMD00061083 | GCA_000011365 | 637000038 | strain page |
| 9234 | | PRJNA714594 | SAMN18312895 | GCA_017565685 | | subspecies page |
| 10154 | | PRJNA714589 | SAMN18312838 | GCA_017565645 | | species page |
| 8857 | | PRJNA224116 | SAMN03459138 | GCF_004114935 | | species page |
| 7915 | | PRJNA224116 | SAMN15581659 | GCF_016031635 | | species page |
| 7297 | | PRJNA497496 | SAMN10259397 | GCA_004114425 | | strain page |
| 9092 | | PRJDB15044 | SAMD00576799 | GCA_030160715 | | species page |
| 7906 | | | | | | |

| | | | | | | | Type Stra |
|---|---|---|---|---|---|---|---|
| **TYGS ID** | **Kind** | **Species cluste** | **Subspecies cluste** | **Preferred name** | **Deposit** | **Authority** | **Other deposits** |
| 104516 | type strain | 1 | 0 | 'Bradyrhizobium xenonodulans | 14ABT | Claassens et al. 2023 | LMG 31415; SARCC 753 |
| 115474 | type strain | 2 | 1 | Bradyrhizobium liaoningense | NBRC 100396 | Xu et al. 1995 | 2281; LMG 18230; CIP 1048 |
| 14517 | type strain | 3 | 3 | 'Bradyrhizobium sacchari' | P9-20 | de Matos et al. 2017 | BR 10280; HAMBI 3667 |
| 15620 | type strain | 4 | 4 | 'Bradyrhizobium forestalis' | INPA54B | Martins da Costa et al. 2 | LMG 10044 |
| 16415 | type strain | 5 | 5 | Bradyrhizobium daqingense | CGMCC 1.10947 | Wang et al. 2013 | LMG 26137; CCBAU 15774; |
| 17297 | type strain | 6 | 6 | Bradyrhizobium cajani | 1010 | Araújo et al. 2017 | AMBPC1010; LMG 29967; C |
| 18506 | type strain | 7 | 7 | Bradyrhizobium huanghuaihae | CGMCC 1.10948 | Zhang et al. 2012 | LMG 26136; CCBAU 23303; |
| 18779 | type strain | 8 | 8 | Bradyrhizobium niftali | CNPSo 3448 | Klepa et al. 2019 | CL 40; U687; USDA 10051 |
| 18997 | type strain | 9 | 9 | Bradyrhizobium shewense | ERR11 | Aserse et al. 2018 | LMG 30162; HAMBI 3532 |
| 2697 | type strain | 10 | 10 | Bradyrhizobium diazoefficiens | USDA 110 | Delamuta et al. 2013 | 3I1B110; ACCC 15034; CNPS |
| 120074 | type strain | 11 | 2 | Bradyrhizobium barranii subsp. | 38S5 | Bromfield et al. 2022 | LMG 31556; HAMBI 3721 |
| 71334 | type strain | 11 | 11 | Bradyrhizobium barranii | 144S4 | Bromfield et al. 2022 | LMG 31552; HAMBI 3722 |
| 8842 | type strain | 12 | 12 | 'Bradyrhizobium zhanjiangense | CCBAU 51778 | Li et al. 2019 | LMG 29279; HAMBI 3648 |
| 92658 | type strain | 13 | 13 | Bradyrhizobium diversitatis | CNPSo 4019T | Klepa et al. 2021 | LMG 31650; WSM 4799 |
| 9288 | type strain | 14 | 14 | Bradyrhizobium vignae | LMG 28791 | Grönemeyer et al. 2016 | 7-2; DSM 100297; NTCCM0 |
| 94558 | type strain | 15 | 15 | Bradyrhizobium iriomotense | NBRC 102520 | Islam et al. 2010 | LMG 24129; EK05 |
| U521176 | user strain | 16 | 16 | DB5 contigs | | | |

ain Genome Server

| Synonymous taxon names | Base pairs | Percent G+C | No. proteins | Goldstamp | Bioproject accessio | Biosample accession | Assembly accessi | IMG OID |
|---|---|---|---|---|---|---|---|---|
| Bradyrhizobium xenonodulans | 8276659 | 6399 | 7625 | | PRJNA680005 | SAMN16869375 | GCA_027594865 | |
| Bradyrhizobium liaoningense | 7830169 | 6395 | 7615 | | PRJDB15044 | SAMD00576800 | GCA_030160735 | |
| Bradyrhizobium sacchari | 8685818 | 6384 | 8163 | | PRJNA224116 | SAMN04849207 | GCF_002068095 | |
| Bradyrhizobium forestalis | 8235644 | 6387 | 7360 | | PRJNA418781 | SAMN08035463 | GCA_002795245 | |
| Bradyrhizobium daqingense | 7885317 | 6374 | 7442 | Gp009388 | PRJNA255603 | SAMN02927912 | GCA_007830205 | |
| Bradyrhizobium cajani | 8372827 | 6396 | 7348 | | PRJNA593773 | SAMN13489115 | GCA_009759665 | |
| Bradyrhizobium huanghuaihai | 9223029 | 6394 | 8622 | Gp009389 | PRJNA255602 | SAMN02927902 | GCA_007830635 | |
| Bradyrhizobium niftali | 9786055 | 6353 | 8664 | | PRJNA529074 | SAMN11254745 | GCA_004571025 | |
| Bradyrhizobium shewense | 9162066 | 6323 | 8018 | Gp010827 | PRJNA224116 | SAMN04487809 | GCF_900094605 | |
| Bradyrhizobium diazoefficiens | 9105828 | 6406 | 8317 | Gp000066 | PRJNA17 | SAMD00061083 | GCA_000011365 | ######## |
| Bradyrhizobium barranii subsp | 10530141 | 6334 | 9234 | | PRJNA714594 | SAMN18312895 | GCA_017565685 | |
| Bradyrhizobium barranii; Brad | 11399526 | 6313 | 10154 | | PRJNA714589 | SAMN18312838 | GCA_017565645 | |
| Bradyrhizobium zhanjiangense | 9342022 | 6294 | 8857 | | PRJNA224116 | SAMN03459138 | GCF_004114935 | |
| Bradyrhizobium diversitatis | 8437014 | 6393 | 7915 | | PRJNA224116 | SAMN15581659 | GCF_016031635 | |
| Bradyrhizobium vignae | 8169430 | 6322 | 7297 | | PRJNA497496 | SAMN10259397 | GCA_004114425 | |
| Bradyrhizobium iriomotense | 9736893 | 629 | 9092 | | PRJDB15044 | SAMD00576799 | GCA_030160715 | |
| | 8460253 | 6319 | 7917 | | | | | |

| |
|---|
| **BacDive** |
| species page |
| strain page |
| species page |
| species page |
| strain page |
| strain page |
| strain page |
| species page |
| species page |
| strain page |
| subspecies page |
| species page |
| species page |
| species page |
| strain page |
| species page |
